# Force-dependent interactions between talin and full-length vinculin

**DOI:** 10.1101/2021.04.26.441533

**Authors:** Yinan Wang, Mingxi Yao, Karen B. Baker, Rosemarie E. Gough, Shimin Le, Benjamin T. Goult, Jie Yan

**Affiliations:** Department of Physics, National University of Singapore, Singapore 117546.; Department of Biomedical Engineering, Southern University of Science and Technology, P. R. China 518055; School of Biosciences, University of Kent, Canterbury CT2 7NJ, UK; Mechanobiology Institute, National University of Singapore, Singapore 117411.

**Keywords:** vinculin, talin, mechanical memory, autoinhibition, magnetic tweezers, mechanotransduction

## Abstract

Talin and vinculin are part of a multi-component system involved in mechanosensing in cell-matrix adhesions. Both exist in auto-inhibited forms, and activation of vinculin requires binding to mechanically activated talin, yet how forces affect talin’s interaction with vinculin has not been investigated. Here by quantifying the force-dependent talin-vinculin interactions and kinetics using single-molecule analysis, we show that mechanical exposure of a single vinculin binding site (VBS) in talin is sufficient to relieve the autoinhibition of vinculin resulting in high-affinity binding. We provide evidence that the vinculin undergoes dynamic fluctuations between an auto-inhibited closed conformation and an open conformation that is stabilized upon binding to the VBS. Furthermore, we discover an additional level of regulation in which the mechanically exposed VBS binds vinculin significantly more tightly than the isolated VBS alone. Molecular dynamics simulations reveal the basis of this new regulatory mechanism, identifying a sensitive force-dependent change in the conformation of an exposed VBS that modulates binding. Together, these results provide a comprehensive understanding of how the interplay between force and autoinhibition provides exquisite complexity within this major mechanosensing axis.

## Introduction

Integrin-mediated adhesions are multi-component molecular complexes that support the physical connection between cells and the extracellular matrix (ECM). At the core of these structures are the transmembrane integrins, *α* - *β* - heterodimers that bind ECM proteins through their large extracellular domains, and are connected to the intracellular actin cytoskeleton via adapter proteins such as talin^1^ and vinculin^2–4^. Integrin adhesions are required for cells to sense the rigidity of their microenvironment, which is important in a variety of processes including tissue formation, maintenance and repair^5, 6^. Hence, understanding the fundamental mechanisms by which integrin adhesions sense and integrate mechanical signals is of crucial importance.

Talin plays a central role in integrin function and mechanosensing. By binding to *β*-integrin tails through its N-terminal four-point-one, ezrin, radixin, moesin (FERM) domain, talin initiates inside-out integrin activation^7, 8^, while its large C-terminal rod domain supports the connection between integrins and F-actin^9^. When talin binds to integrins at one end and to F-actin at the other, it is mechanically stretched due to actomyosin contraction^10–13^. Previous studies have revealed that talin responds to external forces by changing conformation which in turn affects interactions with its binding partners including vinculin^14–20^.

Vinculin, a 116 kDa cytoplasmic protein, has emerged as a key regulator of integrin adhesions^4^. It acts in part by cross-linking integrin-talin complexes to the actin cytoskeleton^4^, and facilitates actin polymerization and nucleation^21^. Hence, vinculin plays a central role in cell adhesion formation, maturation, and turnover^22^. Full-length vinculin (FL-vinculin) is comprised of five domains, D1, D2, D3, D4, which together form the vinculin head that is connected via a proline-rich linker to the vinculin tail domain (Vt) (Fig. 1). Inter-domain interactions within the vinculin head organize the head into a pincer-like structure^23, 24^ (Fig. 1). Vinculin binds to the 11 vinculin binding sites (VBSs) in the talin rod via its D1 domain, and to F-actin through Vt, and to numerous other signaling proteins via interactions with various domains^2, 3, 25, 26^.

**Figure 1.**
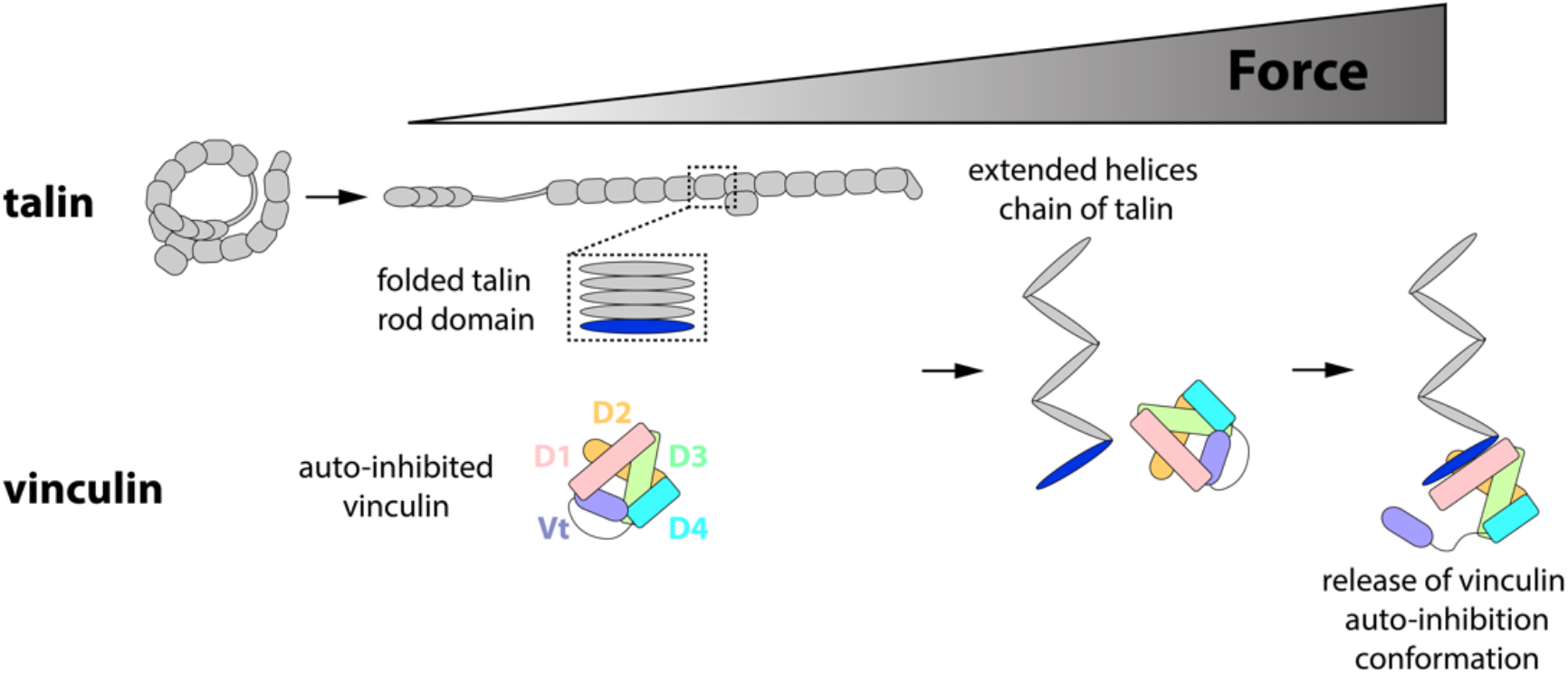
Schematic of the force-dependent vinculin activation by talin. In the absence of force, both the VBS (blue) in talin rod domains and FL-vinculin are autoinhibited. Force is needed to expose the VBS in talin rod domain by unfolding the *α*-helix bundle into an extended chain of *α*-helices. The mechanically exposed VBS can bind to the D1 domain (pink) and compete off the tail, which releases the autoinhibitory conformation.

Vinculin binding to talin requires exposure of the VBS buried in the *α*-helical bundles in the talin rod domains^14–16, 27^. Recent studies have revealed that forces within the physiological range, in the order of several piconewtons (pN), can expose the cryptic VBSs in talin and enable vinculin D1 binding^14–16^. While previous studies have shown that isolated talin VBSs bind vinculin with dissociation constants over a range of 70 - 500 nM^28^, it remains unclear how mechanically exposed VBSs in the context of unfolded rod domains might interact with vinculin. In contrast to an isolated VBS helix, which exists in a force-free environment, the mechanically exposed VBS in talin are under forces of several pN^12, 13, 16^, which may alter the conformation of the VBS helix significantly impacting on the binding affinity^29^ and kinetics^30^. Hence, further studies on the force-dependent conformations of VBSs under a few pN forces and the resulting effects on the vinculin-talin VBS interaction may provide insight into this important linkage.

A further layer of regulation arises from the fact that vinculin is also autoinhibited, and in the absence of other factors, vinculin adopts a compact globular conformation, in which the vinculin head interacts with the vinculin tail suppressing its interactions with most of its binding partners (Fig. 1). As vinculin head binds to its tail with high affinity in vitro^31, 32^, this autoinhibitory interaction is thought to be strong. Several models have been proposed to explain the vinculin activation process at cell adhesions. The widely accepted combinatorial model proposes that at least two binding partners are required to associate with vinculin head and tail simultaneously to overcome the strong head-tail interaction^23, 33^. However, based on the high-affinity interaction between vinculin D1 and isolated talin VBS, the possibility of a single ligand activation model cannot be excluded. Since the vinculin-talin and vinculin autoinhibitory associations are mutually exclusive, the D1-VBS interaction may provide sufficient energy to compete off Vt from D1^28, 34, 35^. In addition, the crystal structure of autoinhibited vinculin shows that autoinhibition is mediated by a number of head-tail interactions, with the D1-Vt interaction being the predominant interaction. As such disruption of the D1-Vt interaction by a talin VBS may destabilize the whole head-Vt interaction, driving a conformational change to a more extended activated conformation^28^.

Another important facet of the dynamics of these linkages is their lifetime. The talin-mediated force-transmission supramolecular linkages have an average lifetime in the order of minutes^36^ although a significant population of talin is immobile in focal adhesions^37^ and at muscle attachment sites^38^. Therefore, the binding of vinculin to talin in cells happens within a limited time window. When talking about the head-tail autoinhibition of vinculin, one should consider whether such autoinhibition can significantly suppress the binding over this physiologically relevant time scale. After binding to talin’s mechanically exposed VBSs, vinculin mediates a cascade of downstream biochemical events through interactions with a plethora of cytoskeletal and signaling proteins^2, 3, 25, 26^. It is reasonable to believe that the longer the activated vinculin is associated with talin, the more persistent the vinculin-mediated mechanotransduction. Hence, the information of the lifetime of vinculin bound to talin is important but has not been investigated in previous studies.

In this study, we show that FL-vinculin can bind to mechanically exposed talin VBS at ~10 nM concentrations, much lower than the dissociation constants measured for isolated VBSs that are not under force^28^. Molecular Dynamics (MD) simulation on isolated VBS reveals a compact helix-hairpin conformation of the talin VBS3 in the absence of tensile force, which autoinhibits the VBS but can be released by physiological forces. The kinetics of the interaction between vinculin and the mechanically exposed VBS is characterized with a fast association rate *k*_*on*_ in the order of 10^6^ *M*^−1^ · *s*^−1^, and a dissociation rate *k*_*off*_ in the order of 10^−2^ *s*^−1^. In addition, by comparing the results with those obtained from vinculin D1, the vinculin head, and an activating vinculin T12 mutant, we determine the influence of inter-domain interactions within vinculin on the kinetics and affinity of the force-dependent vinculin-talin interaction.

## Results

### A force-jump cycle assay to quantitate vinculin-talin complexation at the single-molecule level

The interactions between the vinculin D1 domain and VBS-bearing talin, *α*-catenin and *α*-actinin domains have been extensively studied^14–16, 39, 40^. In contrast, relatively little is known about the force-dependent binding of FL-vinculin to these VBS-bearing mechanosensing proteins. In this study, we wanted to study binding of FL-vinculin to a well characterized mechanosensitive system, and we chose talin domains R4-R6 which contain three *α*-helical bundles and a single VBS buried in R6 (helix 27) (Fig. 2A-B). Talin R4-R6 was tethered between a glass surface and a superparamagnetic bead which enabled force to be exerted onto the domains (Fig. 2A).

**Figure 2.**
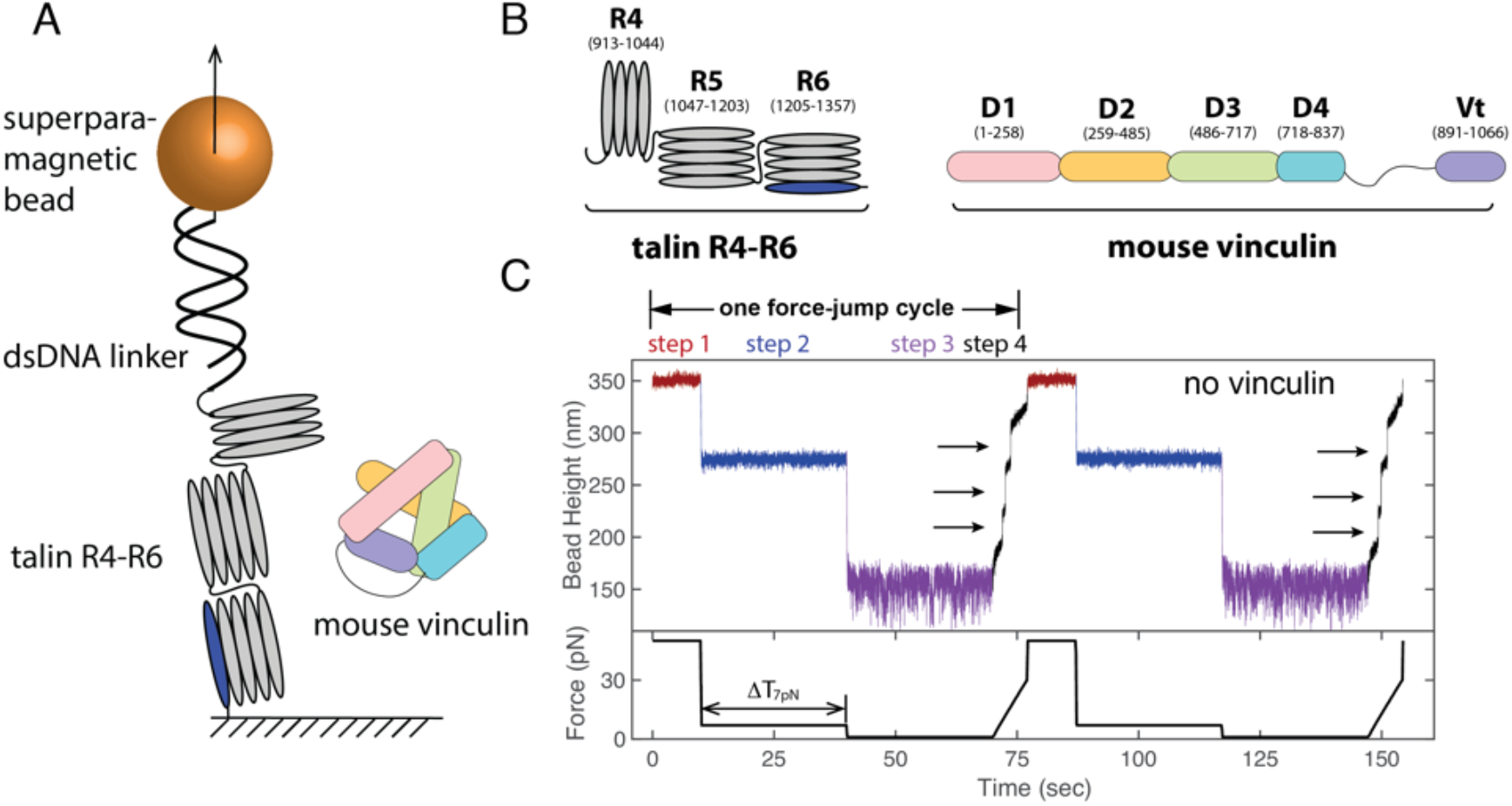
Force-jump cycle to detect and quantify vinculin binding. (A) Schematic of the talin R4-R6 domains tethered between a glass surface and a superparamagnetic bead. A 572-bp DNA linker is added as a spacer. (B) Domain map of talin R4-R6 (left) and FL-vinculin (right). The cryptic VBS in R6 is shown in blue. (C) Force-jump cycle applied to the talin R4-R6 domains in the absence of vinculin. Black arrows indicate the three discrete unfolding steps corresponding to R4-R6 domains. (bottom: the experimental time trace of force change).

To detect and quantify binding of FL-vinculin to mechanically unfolded R4-R6 we implemented a force-jump cycle approach (Fig. 2C). Each force-jump cycle included the following steps: 1) a vinculin displacement step: the single-molecule construct was held at 50 ± 5 pN for 10 seconds to ensure displacement of any bound vinculin within seconds with 100% probability^16^; 2) a vinculin binding step: force-jump to a vinculin-binding force of 7 ± 0.7 pN for a certain time interval, Δ*T*_7pN_ to allow vinculin binding to the mechanically exposed VBS; 3) a talin refolding step: force-jump to domain-folding force of 1 ± 0.1 pN for 30 seconds to allow all domains to refold with 100% probability unless vinculin remains bound to the VBS in talin R6 preventing refolding of R6^16^, and 4) binding detection step: force was increased from 1 ± 0.1 pN to 30 ± 3 pN at a loading rate of 4 ± 0.4 pN/s, during which all the domains unfold. After this step, force was jumped back to 50 ± 5 pN for the next vinculin displacement step, which completes one force cycle.

A vinculin-binding force of ~ 7 pN was chosen to detect the binding of vinculin to mechanically exposed VBS, because it lies within the physiological range of forces (5 – 12 pN) applied to talin in cells^12, 16^. In addition, at this force, refolding has never been observed over a long time scale (> 400 seconds)^16^, which ensured that the VBS was always exposed for binding. At the domain-folding force of ~1 pN, in the absence of other factors, all talin rod domains fold almost immediately. In step 4 for the binding-detection, if no vinculin was bound then three unfolding events would be observed as seen in Fig. 2C. However, if a domain remains unfolded after 30 seconds, this can be attributed to vinculin-binding to the VBS preventing refolding of the VBS-containing R6 domain. Therefore, this assay enables us to monitor whether a vinculin molecule is bound to our talin molecule; if R6 is not able to refold in step 3 due to vinculin remaining bound, only two unfolding events from R4 and R5 would be observed indicating formation of a talin-vinculin complex.

### The response of talin R4-R6 to cyclic force perturbation

Fig. 2C shows a representative time trace of more than 10 independent tethers of bead height change during a force-cycle in the absence of vinculin. At step 1 where the tether was held at 50 ± 5 pN, the bead height fluctuated around a constant average level of 350 nm (data shown in red). The following force-jump to 7 ± 0.7 pN resulted in a large abrupt height decrease to an average level of 274 nm (data shown in blue). The next force-jump to 1 ± 0.1 pN resulted in another abrupt height decrease to an average level of 151 nm (data shown in purple). During the subsequent force-increase scan from 1 ± 0.1 pN to 30 ± 3 pN (data shown in black), three unfolding steps were observed (black arrows), indicating full refolding of all the three domains when the construct was held at 1 ± 0.1 pN in this force-cycle. Here we note that the abrupt bead height changes during sudden large force jumps are contributed from both intrinsic molecular extension change, and bead rotation due to torque rebalance after the force change^41^. In contrast, the stepwise bead height changes during force-increase scans represent molecular extension changes. This is because the force change before and after the steps are less than 0.04 pN, hence the torque remains balanced and therefore the bead rotation is negligible^41^.

### Vinculin binds to mechanically unfolded talin

Repeating the force-cycle in the presence of 10 nM FL-vinculin revealed only two unfolding events (indicated by black arrows) at the binding detection step in the first force-jump cycle (Fig. 3A) with Δ*T*_7pN_ = 30 s. This indicates that one domain did not refold at step 3 when the construct was held at 1 ± 0.1 pN for 30 seconds. Since the failure of the domain to refold was dependent on the presence of FL-vinculin, it was attributed to the vinculin-bound R6. In cycle 2, three unfolding steps were observed, indicating no vinculin was bound to the VBS in the step 2 - vinculin binding step. The probability of having a FL-vinculin bound during any given cycle provides important information on the binding affinity of the interaction.

**Figure 3.**
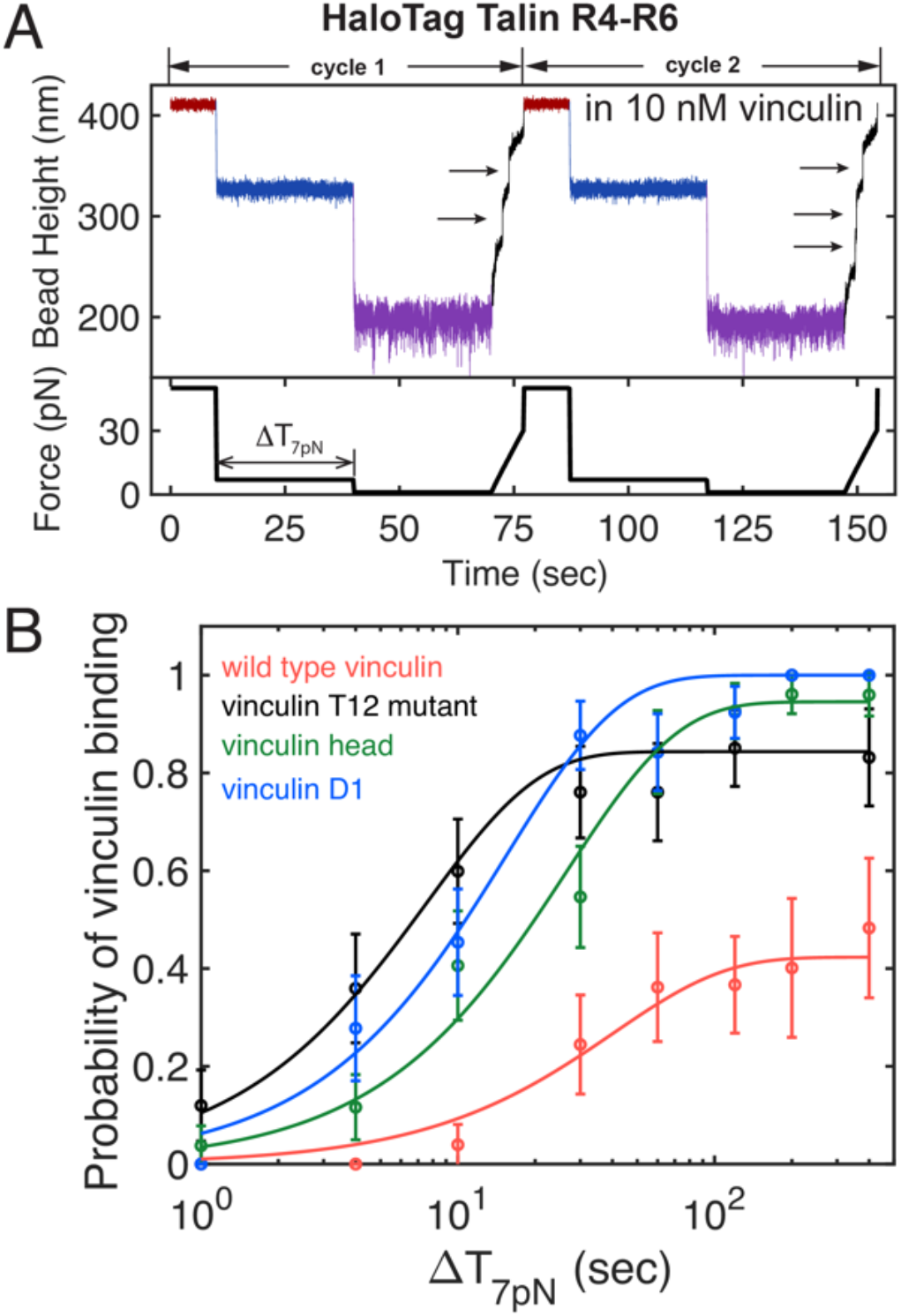
Full-length vinculin binds to mechanically exposed VBS in talin with nM affinity. (A) Representative force-jump cycles applied to detect and quantify vinculin binding. (B) The time evolution of binding probability for wild type vinculin (red), T12 mutant (black), vinculin head (D1-D4, green), and vinculin D1 (blue) are shown. The evolution for vinculin binding probability was taken at Δ*T*_7pN_=1 s, 4 s, 10 s, 30 s, 60 s, 120 s, 200 s, and 400 s.

The fact that the vinculin binding was observed in these experiments indicates that mechanical exposure of the VBS in R6 is sufficient for FL-vinculin binding at nM concentrations, suggesting that the autoinhibitory head-tail interaction of vinculin does is not strong enough to suppress vinculin binding to a mechanically exposed VBS.

### Quantification of vinculin binding to a mechanically exposed VBS

We next sought to quantify the interaction between talin and vinculin by determining the binding kinetics and affinity. To do this, we implemented the force-jump cycle (Fig. 2C) to obtain the probability of FL-vinculin binding to the mechanically exposed VBS in talin R6 at 7 ± 0.7 pN with different holding times Δ*T*_7pN_. At each Δ*T*_7pN_, the binding probability was determined by 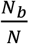, where *N* ≥ 15 was the total force-cycles obtained from multiple tethers and *N*_*b*_ was the number of cycles where vinculin binding was observed. Repeating the force-cycle at different Δ*T*_7pN_, we determined the time evolution of the binding probability *P*(*t*) of FL-vinculin binding to the mechanically exposed VBS in talin R6.

In Fig. 3B, the red data points show the probability of FL-vinculin binding to the mechanically exposed VBS in talin R6 obtained at 10 nM FL-vinculin. Shown in red is the best-fit curve with 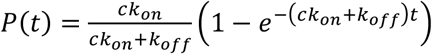, where *c*, *k*_*on*_ and *k*_*off*_ are vinculin concentration, association rate and dissociation rate, respectively. The best-fit parameters are determined to be *k*_*on*_ = 1.0 ± 0.4 × 10^6^ *M*^−1^ *s*^−1^ and *k*_*off*_ = 1.4 ± 0.9 × 10^−2^ *s*^−1^, from which the dissociation constant was calculated to be 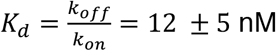. The result indicates that FL-vinculin can directly bind to the mechanically exposed VBS in talin R6 with nM affinity. The standard error was calculated as the standard deviation of means based on bootstrap analysis with 200 repetitions (See Methods). The association rate is in the order of diffusion limited on-rate^42^; hence, the result strongly suggests that the FL-vinculin undergoes a highly dynamic fluctuation between the auto-inhibited closed conformation and an open conformation accessible to the VBS.

### Quantification of vinculin T12 mutant, vinculin head and vinculin D1 binding to mechanically exposed VBS

Having established that FL-vinculin binds to mechanically exposed VBS in talin, we next characterized the interaction in more detail using a series of well-established vinculin constructs. These included the constitutively active “vinculin T12” mutant^31^, the entire vinculin head (D1-D4), and the VBS-binding domain of vinculin (D1) which is expected to bind talin with maximal affinity. Similar binding experiments were performed for each of these vinculin constructs to enable direct comparison with FL-vinculin (Fig. 3B).

#### Vinculin T12 mutant

The vinculin T12 mutant contains four mutated residues in the vinculin tail domain, Vt (D974A, K975A, R976A and R978A)^31^. Previous experiments have shown that the T12 mutant has a weaker head-tail interaction due to disruption of the D4-Vt interface, resulting in stronger binding to talin and enhanced focal adhesion formation and stabilization^31^. In our force-cycle experiments, 10 nM vinculin T12 (Fig. 3B, black curve) bound to the mechanically exposed VBS in talin R4-R6 with the following best-fitting values, *k*_*on*_ = 1.1 ± 0.3 × 10^7^ *M*^−1^ *s*^−1^ and *k*_*off*_ = 2.1 ± 1.1 × 10^−2^ *s*^−1^. The dissociation constant was calculated to be *K*_*d*_ = 1.9 ± 0.5 nM. Compared to wild type vinculin, T12 has a significantly faster association rate but a similar dissociation rate, resulting in a higher binding affinity indicated by a ~6x lower dissociation constant.

#### Vinculin head

The vinculin head comprises D1, D2, D3 and D4 domains that show extensive interdomain interactions, although the construct lacks the autoinhibitory Vt domain. Similar experiments performed in 10 nM vinculin head (Fig. 3B, green curve) gave best-fitting values of *k*_*on*_ = 3.7 ± 0.9 × 10^6^ *M*^−1^*s*^−1^ and *k*_*off*_ = 2.2 ± 1.8 × 10^−3^ *s*^−1^. The dissociation constant was calculated to be *K*_*d*_ = 0.6 ± 0.3 nM. The measured value of *k*_*off*_ is in good agreement with that reported from a previous Fluorescence Recovery After Photobleaching (FRAP) measurement^19^.

#### Vinculin D1

Similar experiments with the vinculin D1 alone (Fig. 3B, blue curve) determined a best-fitting value of *k*_*on*_ = 6.5 ± 1.6 × 10^6^ *M*^−1^*s*^−1^ and a near-zero *k*_*off*_ which cannot be accurately determined by fitting. Together, these results suggest a much higher binding affinity for vinculin D1, which cannot be determined within our experimental time scale, than that of the vinculin head, FL-vinculin and the T12 vinculin mutant.

The best-fitting values of *k*_*on*_ and *k*_*off*_ and the resulting *K*_*d*_ are summarized in Table 1. These results suggest that although all four forms of vinculin can bind the mechanically exposed VBS at nM concentrations, the binding affinity is the highest for D1 and the weakest for FL-vinculin. The dissociation rates for the FL-vinculin and the T12 mutant are similar; therefore, the increased affinity of T12 is mainly caused by a near 10-fold faster association rate compared to the wild type vinculin. This suggests that the dynamic head-tail interaction within the wild type vinculin reduces the time fraction of the open, accessible conformation.

**Table 1.** Kinetic rates and affinity of vinculin binding to mechanically exposed VBS.

|  | FL-vinculin | T12 vinculin | vinculin head | vinculin D1 |
| --- | --- | --- | --- | --- |
| $k_{on} (M^{-1}s^{-1})$ | $1.0 \pm 0.4 \times 10^6$ | $1.1 \pm 0.3 \times 10^7$ | $3.7 \pm 0.9 \times 10^6$ | $6.5 \pm 1.6 \times 10^6$ |
| $k_{off} (s^{-1})$ | $1.4 \pm 0.9 \times 10^{-2}$ | $2.1 \pm 1.1 \times 10^{-2}$ | $2.2 \pm 1.8 \times 10^{-3}$ | not detectable |
| $K_d (nM)$ | $12 \pm 5$ | $1.9 \pm 0.5$ | $0.6 \pm 0.3$ | |

N.B. the $K_d$ of vinculin D1 cannot be obtained via this method as there is no dissociation of the D1 from the Vd1.

The association rates of vinculin head and D1 are faster than that of the FL-vinculin by several folds. In addition, their dissociation rates are ~6-fold (for vinculin head) and much more (for vinculin D1) slower than that of the FL-vinculin. Together, the faster association rates and slower dissociation rates result in the higher affinity of vinculin head and D1 than that of the FL-vinculin.

Overall, these results reveal important differences between the binding of the four forms of vinculin to talin VBS. The most important is that, while the head-tail interaction of vinculin does not inhibit vinculin binding to mechanically exposed VBS, it significantly tunes the affinity mainly via modulating the rates of binding.

### The force-dependent conformations of an exposed talin VBS tune its affinity for vinculin

A striking finding of our study is that the affinity of the talin-vinculin interaction observed under force is significantly higher than the bulk interactions of talin VBS with FL-vinculin measured in solution (70 - 500 nM)^28^. As, the mechanically exposed VBS have enhanced binding affinity relative to isolated VBS in solution, it suggests that forces applied to a talin VBS strongly influence binding to vinculin. The talin-vinculin interaction involves the VBS binding as a helix to the D1 domain via a helix-addition mode of binding (illustrated in Fig. 4A). In the folded talin rod domain, a VBS also adopts a helical conformation as part of the helical bundle. However, an exposed VBS, which is not bound to vinculin, has the potential to adopt various conformations besides the high affinity vinculin-binding helical form. Hence, we hypothesized that the isolated VBS may adopt thermodynamically stable autoinhibited conformations that suppresses its binding to vinculin D1. To evaluate the potential effect of force on VBS conformation we performed 1-*μ*s full-atom molecular dynamics (MD) simulations on a VBS peptide with and without applied forces. The initial VBS conformation used in the simulations was obtained from the crystal structure (PDB 1rkc^34^) of human vinculin D1 bound with VBS3 (i.e., VBS from talin R10), which adopts an *α*-helical conformation in the complex (Fig. 4A).

**Figure 4.**
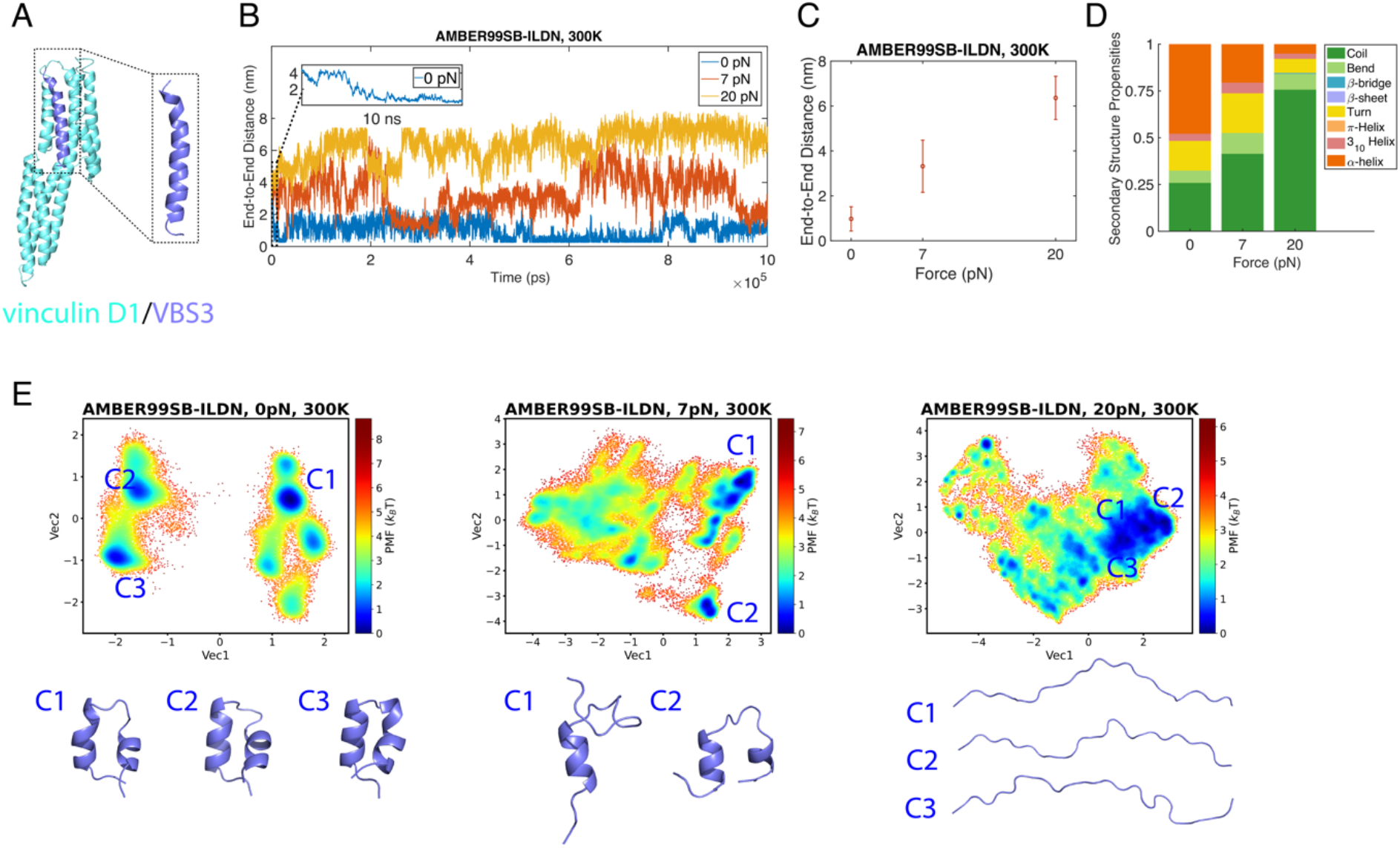
Force-dependent conformations of the talin VBS3. (A) Crystal structure of vinculin D1 in complex with talin VBS3 (PDB 1rkc). The inset shows the *α* -helical conformation of VBS3 used as the initial conformation. (B) Time traces of the end-to-end distance of VBS3 under different tensile forces using AMBER99SB-ILDN force field at 300K. The inset shows the time trace of VBS3 in the first 10 ns in the absence of force. (C) The mean and standard deviation of the end-to-end distance of VBS3 calculated from the time traces at 300K using AMBER99SB-ILDN force field. (D) Secondary structure propensities of VBS3 under different tensile forces using AMBER99SB-ILDN force field at 300K which are defined by DSSP. (E) Free energy landscape as a function of the first two dihedral principal components using AMBER99SB-ILDN force field at 300K at 0, 7, and 20 pN, with the representative conformations of identified local energy minima.

Simulations were performed on the VBS3 by itself in 150 mM NaCl solution starting from the initial *α*-helical conformation derived from the X-ray structure of the VBS3-D1 complex up to 1 *μ*s under AMBER99SB-ILDN^43^ force field, at temperature of 300*K* (See Methods). The time traces of the end-to-end distance of VBS3 show significant dependence on the applied force (Fig. 4B). In the absence of force, the end-to-end distance collapsed from the initial value (~4.4 nm) of the helical conformation to ~ 0.7 nm after 10 ns of simulation (Fig. 4B, inset) and remained in the compact conformation throughout the rest of simulation. At 7 pN, large fluctuations between highly compact conformations and more extended conformations were observed. At 20 pN, the end-to-end distance evolved from the initial 4.4 nm to a larger value (~6.4 nm), indicating transition to conformations that are more extended than the original helical conformation. Consistently, the average extension of VBS3 monotonically increases as the applied force increases (Fig. 4C). Together, the results reveal that forces in physiological range strongly modulate the conformations and the extension of a VBS helix. It is likely that all exposed talin helices exhibit similar force-dependent conformational changes.

To obtain the information on the predominant conformations at each force, we performed the dihedral principal component analysis (dPCA)^44^ and used the first two principal component axes to recast the simulation data (See Methods). Fig. 4E shows the free energy landscape of VBS3 as a function of the first two dihedral principal components of the 1-*μ*s MD simulation in the absence of force, as well as at the forces of 7 and 20 pN at 300*K* using AMBER99SB-ILDN force field, and a few representative predominant snapshots of conformations identified by dPCA at each force. At 0 pN, the free energy landscape represents multiple local energy minima and the corresponding VBS3 conformations all exhibit compact helix-hairpin structures. At 7 pN, the predominant conformations of the VBS3 peptide becomes a mixture of helical and disordered regions. At 20 pN, the VBS3 exists predominantly as a disordered, extended peptide. Fig. 4D summarizes the amino acid secondary structure propensity in 1-*μ*s MD simulation at different forces determined by the Dictionary of Protein Secondary Structure (DSSP) algorithm^45^.

Together, the simulations suggest that the conformation of an isolated talin VBS helix is very sensitive to the force applied over a physiological range. Interestingly, in the absence of forces, an isolated VBS has the propensity to fold into a stable *α*-helical hairpin conformation that not only drastically deforms from the original *α*-helical conformation but also buries several critical residues that mediate the interaction with vinculin D1. This compact form is likely to be a previously unrecognized, autoinhibited conformation that binds D1 with reduced affinity. In addition, at too large a force (e.g., 20 pN), the VBS becomes a completely disordered peptide. At the physiologically relevant forces of a few pN such as 7 pN, the VBS3 is a dynamic mixture of helical and disordered regions which is expected to enhance binding to D1 compared with the stably folded helix-hairpin in the absence of force or the completely disordered peptide conformation at large forces > 20 pN.

Based on the insights provided from the MD simulations, we developed a simple model to understand the force-dependent binding affinity between an isolated talin VBS and the VBS binding D1 domain in vinculin. In this model, three structural states of the VBS are considered (Fig. 5A): the *α*-helical conformation that binds D1 with the highest affinity (state “off, 2”), an autoinhibited *α* -helical hairpin conformation (state “off, 1”) and a disordered peptide conformation that does not bind D1 (state “off, 3”). Besides the three “off” states of VBS unbound by D1, a fourth state where the VBS is bound to D1 (state “on”) exists. Based on these four states, we derived a force-dependent dissociation constant of the VBS-D1 interaction as:

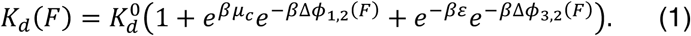

where 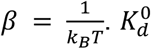 denotes the dissociation constant of the original *α*-helical conformation of VBS binding to D1 in the absence of force. *μ*_*c*_ is the autoinhibitory energy stored in the *α*-helical hairpin which is the free energy difference between the original *α*-helical conformation and the *α*-helical hairpin. *ε* is the free energy difference between the unstructured peptide conformation and *α*-helical conformation of VBS. The value of *ε* tunes the probabilities of the *α*-helical conformation and the unstructured peptide conformation, which can be roughly understood as the stability of the *α*-helical conformation of the VBS. Δ*ϕ*_1,2_(*F*) is the force-induced conformational free energy difference between the “off, 1” and “off, 2” states, and similarly Δ*ϕ*_3,2_,(*F*) is the force-induced conformational free energy difference between the “off, 3” and “off, 2” states. These force-induced conformational free energy differences can be calculated based on the different force-extension curves of the corresponding structural states *i* and *j*, 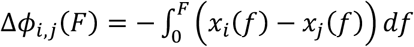, where *x*_*i*_(*f*) is the force-extension curve of the conformation state “off, *i*”. Details of the general physics behind the derivation can be found in our recent publication^29^, and the calculation for this particular case is provided in the supplementary information (Supp. Info. 1).

**Figure 5.**
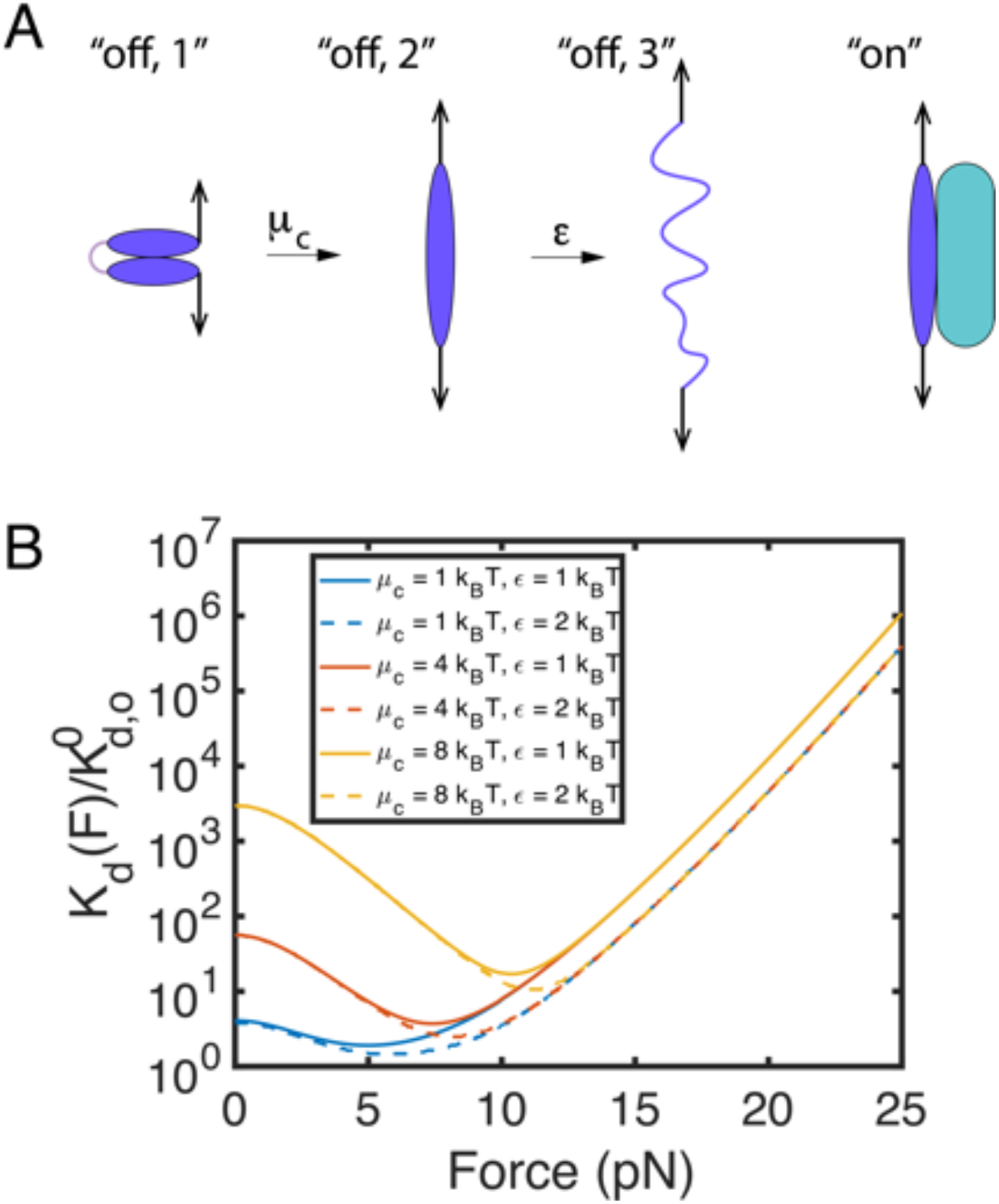
Force-dependency of the VBS conformation and its interaction with VD1. (A) Schematic of four states of an isolated VBS, three unbound, “off” states, and the vinculin D1 bound “on” state. The green rounded rectangle represents vinculin D1 which binds to the helical “on” conformation. (B) Fold change in the force-dependent binding constant, *K*_*d*_(*F*) of the VBS-D1 interaction (Eq. 1), maximal binding affinity is seen in the trough of the curve at 5-11 pN dependent on the value of *μ*_*c*_ and *ε*.

Fig. 5B shows predicted *K*_*d*_ relative to 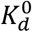 for the VBS binding to the vinculin D1 as a function of force applied to the VBS (Eq. 1). As shown, with a reasonable assumption of *k*_*B*_*T* level of the autoinhibition energy of *μ*_*c*_ in the hairpin-like structure, the equation predicts that forces of a few pN would increase the binding affinity by several times. Furthermore, the force dependent binding constant, has maximal affinity (the trough on the curves in Fig. 5B), that is determined by the stability of the VBS helix, as such helix stability further tunes the talin-vinculin interactions. As a result, further complexity and nuance is added to the talin-vinculin interactions, as even at the level of an exposed VBS the affinity between a VBS helix and vinculin is dynamically regulated by mechanical forces.

## Discussion

The interplay between talin and vinculin dictates mechanotransduction pathways downstream of integrins. A remarkable aspect of the interactions between these two proteins is its complexity. Both talin and vinculin adopt autoinhibited states, and once activated, the lifetimes of their association are largely defined by the mechanical conditions of the system and the history of prior forces that have acted on the linkages. In this study, we investigated the interaction between autoinhibited vinculin and talin, by using a three domain talin construct, R4-R6 that contains a single cryptic VBS in R6. Strikingly, these two proteins do not interact in the absence of force, but mechanical exposure of the VBS in R6 is sufficient for binding to autoinhibited vinculin, revealing that force applied to talin alone is sufficient to activate high-affinity binding of vinculin in the absence of other factors. Furthermore, we identify an additional layer of regulation whereby the affinity of an exposed VBS for vinculin is fine-tuned by force-dependent changes in the conformation of the VBS helix itself. Steered full-atom molecular dynamics simulation reveals that a VBS peptide can adopt an autoinhibited stable helix-hairpin conformation, which can be unfurled at physiological ranges of force. We reason that force may release the autoinhibited helix-hairpin conformation of the VBS enhancing its affinity for vinculin.

Autoinhibition of proteins involved in cell adhesion represents a major mechanism encoding mechanosensitivity^46^ and enabling force-dependent binding constants^29^. Full-length talin and vinculin are both regulated by autoinhibition^23, 24, 47, 48^, and a recent study by Atherton et al. using a mitochondrial targeting assay showed that the two autoinhibited proteins do not interact^49^. In the same study^49^, talin null cells co-expressing full-length vinculin and full-length talin under tension-released condition only formed small peripheral adhesions. However, activation of either protein via mutagenesis was shown to be sufficient to enable their co-localization in focal adhesions. Here, we show that full-length vinculin can bind to the mechanically exposed VBS in talin R6 with nM affinity, indicating that mechanical activation of talin is sufficient to trigger high-affinity binding to full-length vinculin. Together, these results highlight the cascade of activation steps that ultimately lead to the interactions between these two proteins.

Binding of vinculin to a mechanically exposed talin VBS is fast with an association rate of ~ 10^6^ M^−1^ s^−1^, close to the typical diffusion limited association rate^42^. This is surprising, since vinculin was thought to adopt a strongly autoinhibited closed conformation due to the Vt-D1 interaction^23, 24, 31^. The fast association rate observed in our experiment strongly suggests that the autoinhibitory Vt-D1 interaction does not significantly slow down the binding of vinculin to a mechanically exposed talin VBS. Therefore, we propose that vinculin must undergo spontaneous rapid dynamic fluctuation between the open and closed states, making the vinculin D1 domain accessible for rapid binding to a talin VBS.

### Force-independent talin-vinculin interactions enhance the association rate

An unexpected finding of this work is that the vinculin T12 mutant, has the fastest association rate of all of the vinculin constructs tested. The vinculin D1 domain interacts with the mechanically exposed VBS with an association rate of 6.5 ± 1.6 × 10^6^ M^−1^s^−1^, which is faster than that of the entire vinculin head (D1-D4 domains) or wild type vinculin, but slightly slower than the T12 mutant. It is surprising that T12 binds the exposed VBS with a rate faster than either D1 or head, as neither of them contain the autoinhibitory Vt domain. We therefore expected that binding of constructs lacking Vt would be faster than both the wild type vinculin and the T12 mutant. One possible explanation for this result is that the Vt domain and/or the preceding linker region might make some form of non-specific interactions with the unfolded talin domains. Unfolded talin rod domains expose a lot of hydrophobic sidechains, and the helical propensity and hydrogen bond forming tendency of these exposed sequences, means such non-specific interactions are quite possible. Such non-specific interactions would increase the effective local concentration of vinculin, promoting a fast binding rate of the vinculin D1 domain to the VBS. The Vt domain and the linker may therefore mediate pre-complexation of FL-vinculin with partially exposed VBS-containing sites prior to canonical VBS-D1 engagement. This interaction would accelerate binding to the VBS once it is mechanically exposed. Such force-independent talin-vinculin interactions have been proposed previously by Han et al.^50^ and pre-complexation of talin and vinculin was shown to be required for efficient adhesion maturation.

It is also interesting to note that we did not detect spontaneous dissociation of D1 from the mechanically exposed VBS even up to 400 sec for all independently tested, tethered R4-R6 molecules (*N* = 8), suggesting an ultra-slow dissociation rate. The dissociation rate of the head was about 6-fold and 10-fold slower than that of the wild type vinculin and the vinculin T12 mutant, respectively. Therefore, compared with the wild type vinculin and the T12 mutant, the D1 and head domains have the highest affinity mainly due to their slower dissociation rate. This high affinity of D1 locks talin in an unfolded conformation^15^ and expression of vinculin D1 in cells leads to loss of adhesion dynamics^11, 51^ and lethality in flies^52^. Although the vinculin head-tail interaction is insufficient to inhibit vinculin binding to a mechanically exposed talin VBS, our results reveal that it significantly tunes the affinity and binding rates between vinculin and VBS. The T12 mutant with weakened head-tail interaction binds VBS with an affinity 6-fold higher compared to the wild type vinculin, suggesting that the head-tail interaction suppresses the binding of vinculin to a talin VBS.

In this scenario, the master switch for both talin binding to vinculin and for vinculin activation is the mechanical unfolding of talin rod domains. Thus, we conclude that talin is the mechanical switch for talin/vinculin-dependent mechanotransduction. Consistently, previous studies have revealed that talin/vinculin-dependent focal adhesion development and maturation require sufficiently rigid substrate^53^, on which talin is expected to experience considerable mechanical stretching. In addition, earlier work shows that applying external force to focal adhesion sites results in increased recruitment of vinculin to the perturbed sites^54^. These previous results are consistent with the talin mechanical switch model.

### Identification of an additional layer of talin autoinhibition

Talin is regulated by many layers of autoinhibition^55^, from the fully closed form in the cytosol which, upon relief of this head-tail autoinhibition, can open up to reveal the linear arrangement of helical bundles. These bundles are autoinhibited with respect to vinculin binding as they contain cryptic VBS within the bundles themselves. At physiological levels of stretching e.g. 5-15 pN talin bundles unfold into a string of helices, exposing the previously cryptic VBS, a well characterized major mechanosensitive event^15, 16^. These domains remain unfolded even when the force is reduced to just a few pN, providing them with a mechanical memory^16^. Here we define an additional, previously unrecognized, layer of autoinhibition on talin, at the level of the individual VBS. Our MD simulations, and the enhanced affinity for a VBS under force, suggest that forces over a few pN range increase the binding affinity by suppressing the VBS from adopting a low affinity, helix-hairpin conformation. This provides an explanation for the higher binding affinity quantified in our single-molecule experiments, where the VBS is under ~7 pN forces (Table 1), compared with that from bulk measurement where the VBS is not under force^28^. We note that mechanically exposed talin VBSs in live cells are under similar level tensile forces^12, 13^. Further increases in force on a VBS reduce its binding affinity for vinculin by decreasing the *α*-helical fraction of the VBS^15^. Therefore, forces biphasically tune the binding affinity of the exposed VBS for vinculin. This changing, binding affinity as forces on talin fluctuate mean that the affinity for vinculin is dynamically tuned, even for an exposed VBS. Due to all these modulators, the force-dependent interactions of even a single VBS with vinculin are complex, and talin has 11 VBSs.

### Identification of an additional layer of vinculin autoinhibition

One intriguing finding of this work is that D1 binds faster and more tightly to an exposed VBS than the vinculin head, which suggests that the inter-domain interactions within the vinculin head suppress vinculin binding to talin. This raises the possibility of an additional layer of vinculin autoinhibition whereby the D1-VBS interaction is hindered by the other head domains, slowing down the binding rate compared to D1 alone. As vinculin also makes a mechanical linkage when it crosslinks talin to actin, it too will experience mechanical forces acting on it, and these forces, exerted only when vinculin forms a mechanical linkage, will extend vinculin. It is possible that the inter-domain interactions in the vinculin head can be released by force-dependent changes in the conformation of the head enhancing talin binding. Such a scenario would explain our data here, and if this is the case it would suggest that vinculin affinity for talin might also be modulated by force, this time acting on vinculin. Future studies should investigate the effects of forces exerted on vinculin binding to an exposed VBS to confirm this layer of autoinhibition of the talin-vinculin and how force on vinculin modulates its affinity for talin.

### Talin-vinculin complexes as a way to encode mechanical memory

In this study we have focused on the talin module R4-R6 that contains a single VBS in order to work with a simplified system, and we show that force on talin drives talin-vinculin complex formation. Complexation stabilizes the open conformations of the domains and thus alter the lifetimes of the active conformations. Further, stabilization of such interactions in vivo will occur when the vinculin tail engages an actin filament^33^, which both stabilizes the open conformation of vinculin and increases the mechanical linkages on that integrin-talin-actin connection. In a cell, there is the opportunity for incredible diversity and complexity in these mechanical linkages based on the mechanical responses we have identified. Each adhesive structure contains many talin molecules^56^ each of which contains 13 rod domains^27^. The 13 rod domains of talin can be envisaged as binary switches with two states, folded “0” and unfolded “1” and can be converted between these states by changes in mechanical force^16^. Within 8 of the talin rod domains reside 11 VBS, each of which can be exposed by mechanical force to bind vinculin. Therefore, just considering the interaction between vinculin and talin alone, the complexity of the mechanical linkages that can form is staggering. Further complexity emerges with the discovery that the talin switches can be modulated by post-translational modifications such as phosphorylation altering their mechanical response^57^. It is possible that other enzymes may also modify the talin switches in response to signaling, altering the mechanical information stored in these linkages and the resulting signaling hubs that assemble^18^. The patterns of 1s and 0s in each talin molecule will be stabilized by vinculin binding to give persistent mechanical linkages, and the effect of future forces on the mechanical linkages will result in additional exposure of VBS in other domains, explicitly dependent on the talin-vinculin complexes already present. This provides a basis for these mechanical linkages to exhibit mechanical memory as recently described in the MeshCODE theory^58^ with information stored in the shape of these molecules and the cytoskeletal connections that form as a result.

In summary, we provide a comprehensive analysis of the complex interactions between full-length vinculin and a mechanically exposed VBS in talin, defining the fundamental mechanisms that regulate such interactions. In doing so we further expand our understanding of these crucial linkages that control mechanotransduction downstream of integrins.

## Acknowledgements

We thank David Critchley for critical reading of the manuscript. J. Y. was funded by the Singapore Ministry of Education Academic Research Fund Tier 2 (MOE2019-T2-1-099) and the Ministry of Education under the Research Centres of Excellence programme. B.T.G. was funded by BBSRC (BB/N007336/1 and BB/S007245/1). B.T.G. and J.Y. were funded by HFSP (RGP00001/2016).

## Author contributions

Y.W. carried out the single-molecule experiments and molecular dynamics simulations. Y.W., M.Y. and S.L. performed data analysis. Y.W., M.Y., K.B.B., R.E.G. and S.L. contributed to the design and expression of the protein constructs for single-molecule experiments. Y.W., J.Y., and B.T.G. interpreted the experimental and simulation data. J.Y. and B.T.G. supervised the research. Y.W., J.Y. and B.T.G. wrote the paper.

## Methods

### Protein expression and purification

All plasmids were expressed in Escherichia coli BL21(DE3) cultured in Luria-Bertani (LB) media. The stretchable talin R4-R6 fragment was expressed and purified as reported previously^16^. Briefly, the expressed protein was purified via the GST-tag, using glutathione Sepharose resin (GE Healthcare) before being eluted by TEV cleavage. The FL-vinculin and vinculin T12 mutant plasmid constructs were synthesized by GeneArt gene synthesis and cloned into an expression vector (pET-28b). The vinculin head and vinculin D1 were cloned into pET-151 expression vector. The his-tagged vinculin proteins were purified through the his-tag followed by anion exchange using standard protocols^59^. Protein concentrations were determined using the respective extinction coefficients at 280 nm.

### Single-molecule manipulation

An in house-made back-scattered vertical magnetic tweezers was used in the single-molecule manipulation experiments with a spatial resolution of ~1 *nm* and temporal resolution of ~200 *Hz*^41, 60^. Talin R4-R6 domains was tethered to the coverslip through its C-terminal HaloTag/ligand system, while its N-terminus was linked to a superparamagnetic bead through a 572-bp double strand DNA linker. This system was performed in a laminar flow channel. The extension change of the tethered protein was measured based on the height change of the superparamagnetic beads tethered to the protein under force.

The details of the force calibration and control for the single-molecule magnetic tweezers experiments have been described in previous papers^41, 60^.

### Determination of *k*_*on*_, *k*_*off*_, *K*_*d*_ and error estimation

The binding kinetics involve the association and dissociation of binding which are characterized by the association rate *k*_*on*_ and the dissociation rate *k*_*off*_, respectively. Denoting *P* as the probability of the VBS in the unfolded talin R6 bound by vinculin, it satisfies the equation: 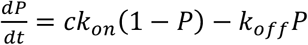, where *c* represents the vinculin concentration. With the well-controlled initial condition *P*(0) = 0, which refers to the assured unbound condition of talin R6 VBS at the starting point of step 2 (Fig. 2C), the equation can be solved as 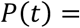 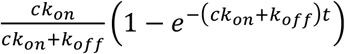.

By implementing the force-jump cycles, the cycles with vinculin binding and those without vinculin binding at the vinculin-binding force (7 pN) can be recorded. At each time interval at 7 pN, Δ*T*_7pN_, an array *A* comprising *N* elements of 0 or 1 was generated, where *N* ≥ 15 is the total number of cycles from multiple independent tethers. Elements of “0” and “1” indicate the cycles where the VBS was “unbound” and “bound”, respectively. In our experiments, the force cycles were performed at the following time intervals Δ*T*_7pN_ = 1 s, 4 s, 10 s, 30 s, 60 s, 120 s, 200 s, 400 s.

After that, bootstrap analysis was performed for 200 repetitions to estimate the mean and the error of the fitted rates. Each bootstrap analysis randomly chooses *N* data points from the array *A* with replacement^61^ and calculated the mean value of the *N* randomly selected data points, which is probability of binding *P*_*i*_(Δ*T*_7pN_), at each Δ*T*_7pN_, where *i* = 1, ···, 200 refers to the *i*^*th*^ bootstrap analysis. For each bootstrap analysis, the resulting *P*_*i*_(Δ*T*_7pN_) was fitted with the function of 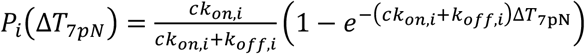, from which the best-fitting values of *k*_*on,i*_ and *k*_*off,i*_ were obtained. Based on the fitted values of association rate *k*_*on,i*_ and dissociation rate *k*_*off,i*_, the dissociation constant *K*_*d,i*_ can thus be determined by 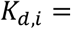 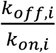. Upon completion of 200-repetition bootstrap (i.e., *i* was taken from 1 to 200), the mean values of the association rate *k*_*on*_, dissociation rate *k*_*off*_, and dissociation constant *K*_*d*_ were determined as the average over all *k*_*on,i*_, *k*_*off,i*_, and *K*_*d,i*_, respectively. The standard error associated with *k*_*on*_, *k*_*off*_, and *K*_*d*_ were determined as the standard deviations of *k*_*on,i*_, *k*_*off,i*_, and *K*_*d,i*_, respectively.

### Molecular dynamics (MD) simulation

The MD simulation was performed using GROMACS 2020.2^62, 63^. The initial talin VBS3 (YTKKELIESARKVSEKVSHVLAALQA) structure was taken from the X-ray structure of the VBS3-human vinculin D1 complex (PDB 1rkc^34^), which adopts the *α*-helical conformation. The simulations were performed under AMBER99SB-ILDN force field^43^ using TIP3P^64^ water model. The initial VBS3 molecule was immersed in periodic cuboid water box filled with 0.15 M NaCl solution. A cutoff distance of 1 nm was applied to the Lennard-Jones interactions and short-range electrostatic interactions. Long-range electrostatic interactions were calculated using Particle-Mesh Ewald (PME) method with a grid spacing of 0.16 nm and 4^th^ order interpolation.

500 steps of steepest descent energy minimization were performed to the simulation system to ensure a reasonable starting structure. Thereafter the energetically minimized system was subjected to a 100-ps NVT equilibration heating and stabilizing the system at 300 K; followed by a 100-ps NPT equilibration stabilizing the pressure of the system. Upon completion of energy minimization and two-step equilibration, 1-μs MD simulation was performed during which the system coordinates were stored every 10 ps for further analysis.

To apply constant force (7 pN and 20 pN) to the VBS3 molecule in MD simulation, the N-terminal tyrosine residue was fixed and C-terminal alanine residue was subject to the corresponding constant force.

### Dihedral angle principal component analysis (dPCA) and free energy landscape

Principal component analysis (PCA) is a dimensionality-reduction method used to identify and retain the most important degrees of freedom of a dynamic simulation system^65, 66^. dPCA has been developed to use the sine and cosine transformed backbone dihedral angles as internal coordinates in the PCA of the MD simulations^44^.

In this work, 48 peptide backbone dihedral angles of the 26-aa VBS3 peptide were used to perform the dPCA. Upon extraction of the dihedral angles from simulation trajectory and implementation of sine and cosine transformation of the 48 dihedral angles, covariance matrix can be calculated based on the sine and cosine variables obtained from the trajectory of dihedral angles. By diagonalizing the covariance matrix, eigenvectors *Vec*_*i*_ and eigenvalues *λ*_*i*_ can be obtained and organized in an eigenvalue-descending order, which means *λ*_*i*_ represents the largest eigenvalue. Thereafter in this work, the first two eigenvectors *Vec*_1_ and *Vec*_2_ associated with the first two largest eigenvalues (i.e., the first two principal components) were chosen to recast the simulation data by projecting the data onto *Vec*_1_ and *Vec*_2_.

Subsequently, the free energy landscape along the first two principal components can be expressed by 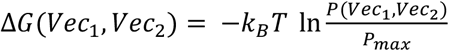, where *k*_*B*_ is the Boltzmann constant, *T* is the temperature, *P*(*Vec*_1_, *Vec*_2_) refers to the probability distribution of the system around (*Vec*_1_, *Vec*_2_), and *P*_*max*_ is the maximum value of the probability distribution.

### Protein secondary structure assignments by DSSP (Dictionary of Protein Secondary Structure)

DSSP is an algorithm assigning secondary structure to the protein residues on the basis of hydrogen bond patterns^45^. The DSSP defines 8 types of secondary structures: a-helix, 3_10_ helix, *π*-helix, hydrogen bonded turn, b-sheet, b-bridge, bend, and coil. In this work, the secondary structure of VBS3 residues were analyzed by using GROMACS do_dssp command with calling the dssp program.

