## Supplementary Information for "Force-dependent interactions between talin and full-length vinculin"

### Supp. Info. 1. Derivation of force-dependent binding affinity of D1-VBS interaction based on a four-state model

Details of the general physics behind the derivation can be found in our previous publication [1], below the calculation for this particular case is provided.

At equilibrium, the dissociation constant $K_d$ can be expressed by the ratio of bound state probability to the unbound probability by $\frac{c}{K_d} = \frac{p_{on}}{p_{off}}$, where $c$ is the concentration of free ligand molecules. In this four-state model, on the basis of the Boltzmann distribution of the states and the definition of the dissociation constant, the following relation holds:

$$\frac{c}{K_d(F)} = \frac{p_{on}(F)}{p_{off}(F)} = \frac{e^{-\beta g_{on}(F)}}{e^{-\beta g_{off,1}(F)} + e^{-\beta g_{off,2}(F)} + e^{-\beta g_{off,3}(F)}},$$

where $\beta = \frac{1}{k_B T}$ and $g_i(F)$ refers to the free energy of state "$i$". Based on the equation, below derivation follows:

$$\frac{c}{K_d(F)} = \frac{e^{-\beta\,[\,g_{on}(F) - g_{off,2}(F)\,]}}{1 + e^{-\beta\,[\,g_{off,1}(F) - g_{off,2}(F)\,]} + e^{-\beta\,[\,g_{off,3}(F) - g_{off,2}(F)\,]}}$$

$$= \frac{e^{\beta \Delta g_0}}{1 + e^{\beta \mu_c} e^{-\beta \Delta\phi_{1,2}(F)} + e^{-\beta\varepsilon} e^{-\beta \Delta\phi_{3,2}(F)}}$$

where $\Delta g_0$ is the binding energy of vinculin to the exposed and valid binding site of the target molecule (i.e., state "off, 2") at zero force, $\mu_c$ is the free energy difference between the $\alpha$-helical conformation (state "off, 2") and the $\alpha$-helical hairpin (state "off, 1"), $\varepsilon$ is the free energy difference between the unstructured peptide conformation (state "off, 3") and $\alpha$-helical conformation (state "off, 2") of VBS. $\Delta\phi_{i,j}(F) =$

$-\int_0^F \left(x_i(f) - x_j(f)\right) df$ is the force-induced conformational free energy difference between the "off, $i$" and "off, $j$" states, where $x_i(f)$ is the force-extension curve of the structural state "off, $i$". By denoting $K_d^0 = ce^{-\beta\Delta g_0}$ which represents the zero-force dissociation constant of vinculin to the exposed and valid binding site, force-dependent dissociation constant $K_d(F)$ can be expressed by

$$K_d(F) = K_d^0\left(1 + e^{\beta\mu_c}e^{-\beta\Delta\phi_{1,2}(F)} + e^{-\beta\varepsilon}e^{-\beta\Delta\phi_{3,2}(F)}\right). \quad (1)$$

Box 1 summarizes the models and parameters of force-extension curves of VBS3 used to predict the value of $K_d(F)/K_d^0$ in Fig. 5B.

| Box 1 | **Model and parameters of force-extension curves of VBS3** |
|---|---|

**$\alpha$-helical hairpin** (state "off, 1") – described by the rigid body with a size of $L_{hh}{\sim}1\,nm$ which is the mean value of zero-force end-to-end distance of VBS3 obtained by the MD simulation (Fig. 4C). Its force-extension curve is $x_1(f) = L_{hh}[\coth\left(\frac{fL_{hh}}{k_BT}\right) - \frac{k_BT}{fL_{hh}}]$.

**$\alpha$-helix** (state "off, 2") – described by the rigid body with a size of $L_{ah,VBS} = N_{VBS}l_{ah,aa}$, where $N_{VBS} = 26$ is the number of amino acids in VBS3 [2] and $l_{ah,aa} = 0.15\,nm$ is the extension per amino acid in the form of $\alpha$-helix [3, 4]. Its force-extension curve is $x_2(f) = L_{ah,VBS}[\coth\left(\frac{fL_{ah,VBS}}{k_BT}\right) - \frac{k_BT}{fL_{ah,VBS}}]$.

**unstructured peptide chain** (state "off, 3") – described by WLC model where the peptide chain has a persistence length of $A_{pt}{\sim}0.8\,nm$ [5] and a contour length of $L_{pp} = N_{VBS}l_{aa}$, where $N_{VBS} = 26$ and $l_{aa}{\sim}0.38\,nm$ is the contour length per amino acid [6]. Based on the WLC model, the force-extension curve of VBS3 peptide $x_3(f)$ can be obtained by solving the inverse function of the Marko-Siggia formula [7]: $\frac{fA_{pt}}{k_BT} = \frac{x_3}{L_{pp}} + \frac{1}{4(1-x_3/L_{pp})^2} - \frac{1}{4}$.
